# Energy use efficiency may mediate metabolic thermal adaptation in *Daphnia magna*

**DOI:** 10.1101/2025.08.01.668087

**Authors:** Thomas Ruiz, Martin J. Kainz

## Abstract

Thermal adaptation in ectotherms is a critical response to global warming. Resting metabolic rate (RMR), reflecting the energy required for body maintenance, is a key factor of thermal adaptation due to its fundamental role in regulating energy acquisition and allocation within other biological functions. Yet, the importance of thermal adaptation of RMR in controlling individual response to warming still requires to be elucidated. Here, we investigated physiological mechanisms underlying thermal adaptation in two *Daphnia magna* lines with contrasting thermal preferences, focusing on RMR but also energy use efficiency (EUE) over a large thermal gradient. Using structural equation modelling we showed that adaptation of EUE between the cold and warm lines was able to mitigate the consequences of RMR thermal variability on individuals’ growth rate. These findings highlight the pivotal role of EUE in ectotherm thermal adaptation, offering key insights to predict the metabolic response of these ectotherms to climate change. Under a context of global warming, ectotherms metabolic response is critical as energy balance and allocation within individuals have repercussions on energy transfers within food web. Accurate predictions of metabolic thermal response of ectotherms may thus constitute an important step toward predicting ecosystem functioning under warming.

## Introduction

Given the accelerating pace at which global change occurs, understanding how organisms, and especially ectotherms, adapt to warmer environments has become a timely challenge in ecology (Rodgers 2021; Jørgensen et al. 2022). The thermal performance of ectotherms is fundamentally driven by metabolic rate, a first-order physiological process directly linked to energy acquisition, allocation, and expenditure (Brown et al. 2004; Kooijman 2009). Temperature has a direct bearing on metabolic rate; as temperature rises, the metabolic rate initially increases due to thermodynamic enzyme kinetics, peaks at a physiologically optimal temperature, and subsequently declines due to oxygen limitations, membrane instability, and nervous and/or mitochondrial dysfunction (Pörtner 2010; Divakaruni and Brand 2011; Wang et al. 2014; Ern et al. 2015). This thermal-response pattern, known as thermal performance curve (TPC), resonates on other life-history traits such as somatic growth or reproductive rates. Such thermal performance curves delineate species’ thermal niches and their potential acclimation capacities to environmental change (Angilletta 2006).

The TPC framework is typically characterised by three key parameters: maximum performance level, thermal optimum, and performance breadth. The maximum performance level reflects the peak functionality of an organism setting the maximum fitness level. The thermal optimum identifies the specific temperature at which this peak performance occurs, providing insights into an organism’s preferred thermal habitat. Finally, performance breadth captures the range of temperatures over which an organism can maintain functional performance, providing a measure of thermal flexibility or tolerance (Knies et al. 2006; see **Fig.1A**). Taken together, these parameters define the limits and potential of individuals to survive and thrive under varying thermal regimes (Angilletta 2004, 2006). Individuals with broader performance breadths may be more resilient to environmental fluctuations, while those with higher maximum performance levels at specific thermal optima might be more competitive in stable, narrowly defined niches (Angilletta et al. 2010). Comparisons of TPC parameters among individuals or populations enable to predict responses and competitiveness under environmental changes (Knies et al. 2006). For example, thermal adaptation may lead to shifts of thermal optima to lower or higher temperatures or induce trade-offs between performance breadth and maximum performance level (Knies et al. 2009; Angilletta et al. 2010). These comparisons are essential for modelling population/species distributions and anticipate their response under ongoing climate change.

**Figure 1.**
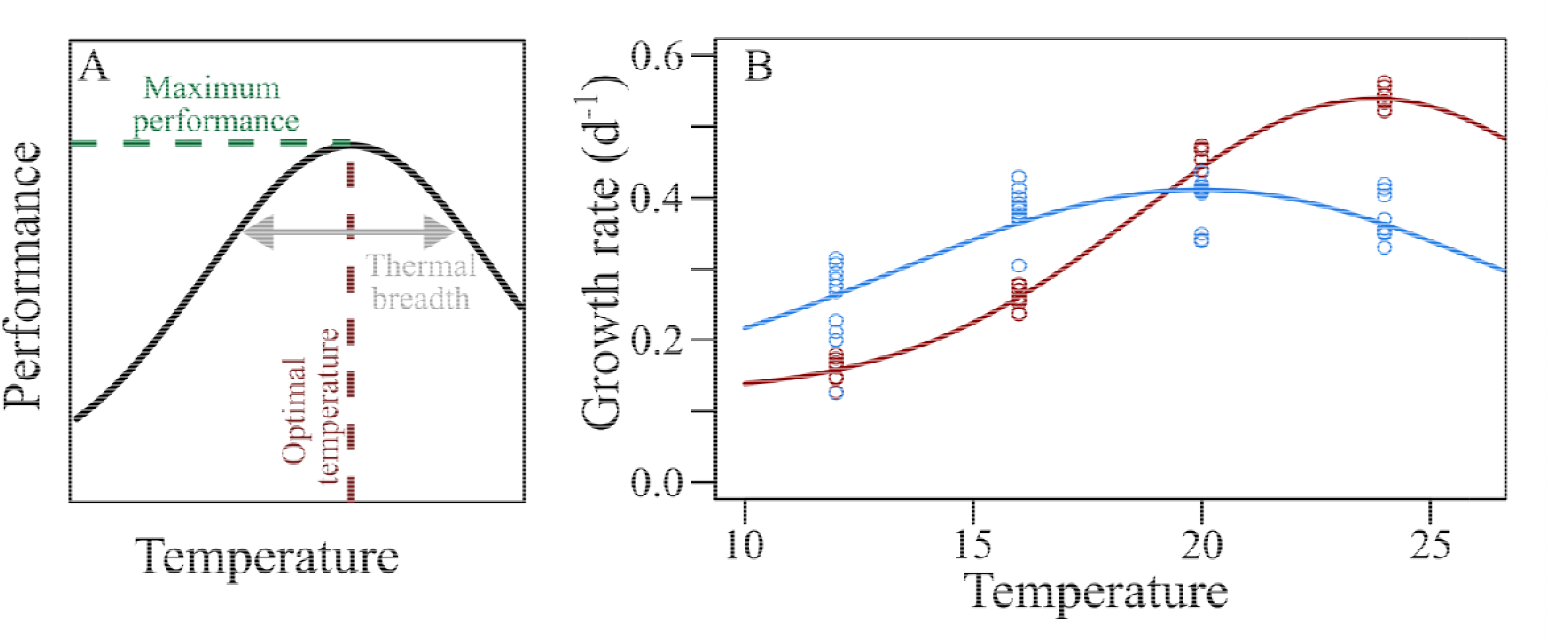
Theoretical representation of a thermal performance curve (TPC) highlighting key parameters (A) and TPC for growth rate for the cold (blue) and warm (red) clonal lines (B). Cold line exhibit a lower maximum performance, a lower optimal temperature but a larger thermal breadth than the warm line.

While TPCs have advanced our understanding of ectotherm thermal adaptation, significant gaps remain in deciphering how resting metabolic level mediates individual thermal responses with respect to performance. Recent studies suggest that populations adapted to different thermal regimes should exhibit distinct metabolic strategies, subsequently affecting energy budgeting and thermal performance (Mesas and Castañeda 2022; Roberts and Williams 2022). For instance, departing from the concept of “counter-gradient variation” (Conover and Schultz 1995) and Krogh’s rule of cold adaptation (Ege and Krogh 1914), several authors posit that warm-adapted individuals may exhibit lower resting metabolic rate (RMR; i.e., the minimal energy required for maintenance) than cold-adapted individuals when exposed to the same temperature. This strategy is expected to reduce the energetic costs for maintenance at higher temperature, thus enabling greater resource allocation to growth or reproduction (Seebacher et al. 2015). However, empirical evidence is contradictory, with some studies supporting metabolic compensation in warm environments (Seebacher et al., 2015; Einum et al., 2019; Jutfelt, 2020), while others reporting minimal or no adaptation of RMR to thermal conditions (Alton et al. 2024). Moreover, the causal relationship between RMR and growth performance remains debated (Arnold et al. 2021), even more under a changing thermal context. Higher RMR could reflect an overall increase of energetic capacity or an energy trade-off from somatic growth to maintenance (Nilsson 2002; Burton et al. 2011; Careau and Garland 2012; Ruiz et al. 2021). Such trade-off between growth and maintenance, and more precisely the ratio between maintenance and growth can be seen as an energy use efficiency (EUE) as it reflects the ability of organism to allocate energy for growth processes rather than to the maintenance of basic body functioning. Disentangling the relative contributions of RMR level and EUE in mediating somatic growth under varying thermal conditions appears critical to adequately define ectotherms thermal adaptations capacities.

To tackle this problem, we examined the interplay between resting metabolic rate and growth performance over a wide temperature gradient in two lines of the ectotherm and aquatic key grazer, *Daphnia magna,* with contrasting thermal adaptation. By combining controlled experiments with a structural modelling approach, this work aims to elucidate the causal and mechanistic links between temperature, thermal adaptation, resting metabolic rate, and growth performances. We specifically hypothesized that, beside the direct consequences of thermal adaptation of RMR, the growth performance of individuals under a changing thermal context may be mediated by the efficiency of energy use within organisms so that energy use efficiency may constitute a key factor for ectotherm thermal adaptation.

## Materials and method

### Organism’s sampling and maintenance

*Daphnia* populations were sampled at two different locations. One population was sampled from a large pond in northern Austria (48.756762; 15.000829) set in a cold climate (10°C when sampled), the other population from a small pond in France (45.646754, 2.458708) located in a warmer climate (26°C when sampled). At the exception of thermal differences, both ponds have a comparable trophic status (i.e., eutrophic). In each population, we randomly sampled 5 individuals of *D. magna* and kept them at 16°C and 24°C, respectively, for the Austrian and French *Daphnia* populations, during 4 generations. Each individual of each population has been kept isolated and clonal lines have been bred independently. At the beginning of the experiment, we selected the clonal line with the highest performance for each group (Austrian and French populations). By doing so, we aimed at getting *D. magna* clonal lines with different thermal history, one line being cold-adapted (i.e. Austrian line), and the other one being warm-adapted (i.e. French line). During the maintenance period, individuals were kept in ADaM medium with the green alga *Chlamydomonas reinhardtii* (3 mg C L^−1^) as food source and under a 12:12 day:night cycle. Prior to the experiment, one individual of each population has been isolated and kept at least for 3 generations to select two pure clonal lines with different thermal preferences.

### Experimental protocol

At the beginning of the experiment, both *Daphnia* lines were acclimated to 20°C for one month to focus on constitutive differences between clonal lines (evolutionary adaptation) rather than short-time acclimation processes. After this month, neonates (less that 12h old) from the third clutch of parent-individuals isolated from the selected clonal line were used in the experiment. At the beginning of the experiment, individuals of each clonal lines were randomly distributed (n=10 each) into 250mL glass jars at 4 temperature treatments (12°C, 16°C, 20°C, and 24°C) and in strictly controlled conditions. In addition, a subsample of neonates from each clonal line was used to determine initial dry weight. To ensure stability of the water composition during the experiment, individuals were kept in daily renewed Volvic^©^ water (Ruiz et al. 2022) supplemented with *C. reinhardtii* (3 mg C L^−1^) as food source and under a 12:12 day:night cycle. Individuals of each treatment were maintained in these conditions until reaching maturity (i.e. egg deposition in brood pouch) or alternatively until 10 days in the absence of egg production. At the end of the experiment, individuals were individually submitted to metabolic rate measurements using respirometry (see below). Subsequently, individuals were dried for 48 h at 50°C to measure their dry weight and calculate juvenile growth rate as a proxy for population growth (Lampert and Trubetskova 1996).

### Metabolic rate measurement

Prior to respirometry measurements of metabolic rate, daphnids were fasting for 6 hrs. This method has been shown to minimize the digestive activity as well as energetic cost dedicated to somatic growth, thus matching as closely as possible their RMR (Ruiz et al. 2018). Daphnids were subsequently placed individually into 24-well microplates with one oxygen sensor spot at the bottom of each 200 µL well (SDR SensorDish ®; PreSens Precision Sensing) filled with Volvic^©^ water and sealed with adhesive film. The system was placed into temperature-controlled water baths to be maintained at experimental temperature during the entire measurement. To correct *Daphnia* oxygen consumption by baseline physical oxygen diffusion we simultaneously performed blank measurements, which also permitted to confirm the thermal equilibrium of the system. Oxygen concentrations were monitored during a maximum of 2 hrs to avoid both oxygen limitation and carbon dioxide accumulation during the measurements. Additionally, we verified the linearity of the oxygen decline rate during measurements and any deviation from linearity, especially at low oxygen level potentially highlighting hypoxic stress, has been removed from data analysis. Oxygen concentrations in each well were integrated over the linear section of the oxygen curve and reported to individual mass to estimate mass-specific individual oxygen consumption.

### Model and data analysis

To test the effects of clonal line and temperature on growth rate, we fitted thermal performance curves through nonlinear least squared regression following the modified Gaussian distribution as follows:

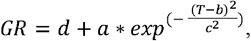

where *GR* is the juvenile growth rate, *d* the minimal value reported for the growth rate in the dataset, and *T* the experimental temperatures. The parameters *a, b,* and *c* were calculated by the model. Parameter *a* reflects the difference between minimum (*d*) and maximum values, so that (*a* + *d*) reflect the maximal growth rate and constitute a proxy for overall performance. Parameter *b* is the optimal temperature. Parameter *c* defines the shape of the curve around the optimum. In addition, we defined the curve breadth as the thermal range (in °C) in which the somatic growth rate remains >90 % of its maximum. Non-overlapping 95% confidence interval (hereafter: CI-95%), generated by bootstrapping approaches, were used to define significant differences in these parameters depending on the clonal line. Experimental values for juvenile growth rate were calculated as:

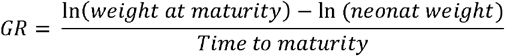

Oxygen consumption, used as proxy for RMR, was related to individual body mass and the interaction with temperature and clonal line was tested using ANOVA after linearization through log transformation. In addition, RMR was corrected for body mass using a scaling coefficient of 0.7 matching previous observation of RMR-mass relation in juvenile *D. magna* fed *Chlamydomonas sp.* (Ruiz et al. 2018, 2021) to obtain a mass-corrected RMR (mcRMR) allowing for comparisons regardless of individual body mass. The thermal dependency of mcRMR, depending on clonal lines, was tested with a similar non-linear least square method as described for the somatic growth rate above.

We defined the ratio between total oxygen consumption from birth to maturity reported versus body mass at maturity; i.e., the amount of body tissues produced per unit oxygen consumed. This ratio reflects the direct link between individual resting metabolic level and biomass production and thus constitutes a proxy for estimating the energy use efficiency (EUE), which can be seen as the balance between energy dedicated to maintenance versus growth. We therefore modelled the body mass development of individuals over time at an hourly interval from birth to maturity (i.e. onset of egg production), departing from measured growth rates, and used it to integrate the instantaneous oxygen consumption over the duration of the experiment. The energy use efficiency is the ratio between total body mass production and total oxygen consumption. Differences of the EUE thermal response were tested using ANOVA for each clonal line independently as the thermal response of EUE in our thermal range was nearly linear and matched the assumption for normality and homoscedasticity.

To define how the mcRMR and EUE underpinned the thermal response of individual growth rates, we developed an incremental structural equation model that accounted for the non-linearity of the mcRMR thermal response. We used temperature as an exogenous variable and modelled the response through nonlinear least square for mcRMR and EUE on our thermal range. For homogeneity, EUE was modelled following a similar approach as mcRMR despite its near-linearity, as the goodness of fit was not different between linear or non-linear regression. The comparison between predicted versus measured values was performed through ANOVA to define the proportion of variance explained by temperature variation as well as its significance and a partial omega squared was used as effect sizes of temperature on both endogenous variables (mcRMR and EUE). We then incremented the model up to individual growth by integrating the linear relation linking mcRMR and EUE to growth rate. Here, we directly estimated the significance and effect size (ω^2^) from ANOVA results. We predicted growth rate following this linear model departing from the mcRMR and EUE values predicted by the nonlinear least squared as established above. Growth rate was thus modelled from temperature through its effect on mcRMR and EUE. The overall accuracy of the model was tested by comparing predicted versus measured values of growth rate by F statistic testing the H_0_ hypothesis of non-independence of predicted versus measured values. We additionally defined the root mean square error of the model (RMSE) as an indicator of its fitting accuracy. Values close to 0 reflect a good fit and the threshold value is generally considered at 0.05. All model and analysis were performed using R software (R-core team).

## Results

### Growth rate

Thermal performance curves for somatic growth rates exhibited significantly different shapes between the two clonal lines. The thermal optimum for growth was 4°C lower for the cold-adapted line (19.9°C [CI-95%: 19.5; 20.7]) compared to the warm-adapted line (23.8°C [CI-95%: 23.1; 24.9]), highlighting their adaptation to distinct thermal environments. Despite having a significantly lower maximal growth rate of 0.42 [CI-95% 0.33; 0.44], the cold line had a tolerance breadth of 10.9°C [CI-95% 9.2; 12.9], a value increasing by 36% in comparison to the warm line showing a tolerance breadth of 8.0°C [CI-95% 7.1; 9.1], but a higher maximal growth rate of 0.55 [CI-95% 0.52; 0.56] (***Fig.1B, Table S1***).

### Resting metabolic rate

Across both clonal lines, body mass (F_(1;76)_=578.7; p=1.44e^−35^) and exposure temperature (F_(1;76)_=11.0; p=1.43e^−3^) had the strongest effect on individual RMR, consistent with established metabolic theories (Gillooly et al. 2001; Brown et al. 2004; Kooijman 2009). The direct effect of the clonal lines on RMR thermal response, excluding interaction with body mass, was only marginal (F_(1;76)_=2.33; p=0.13). The comparison of the TPC for mcRMR showed that, while thermal response pattern remained similar between clonal lines (ANOVA: F_(1;76)_=2.33; p=0.13), the warm line exhibited a significantly higher RMR than the cold line at each tested temperature, reaching maximal values of 9.5 ng O_2_ µg DW h^−1^ [CI-95%: 8.9; 11.5] and 7.14 ng O_2_ µg DW h^−1^ [CI-95%: 6.85; 7.55], respectively (***Fig.2-A***). The tendency of a lower thermal optimum (20.24°C [CI-95%: 19.1; 22.65]) and larger thermal breadth (16.2°C [CI-95%: 10.4; 19.3]) for the cold versus warm line (optimum: 23.04°C [CI-95%: 19.4; 44.8]; breadth: 10.9°C [CI-95%: 8.9; 14.4]) appears marginal for RMR conversely to growth rate (***Fig.2-A, Table S2***).

**Figure 2.**
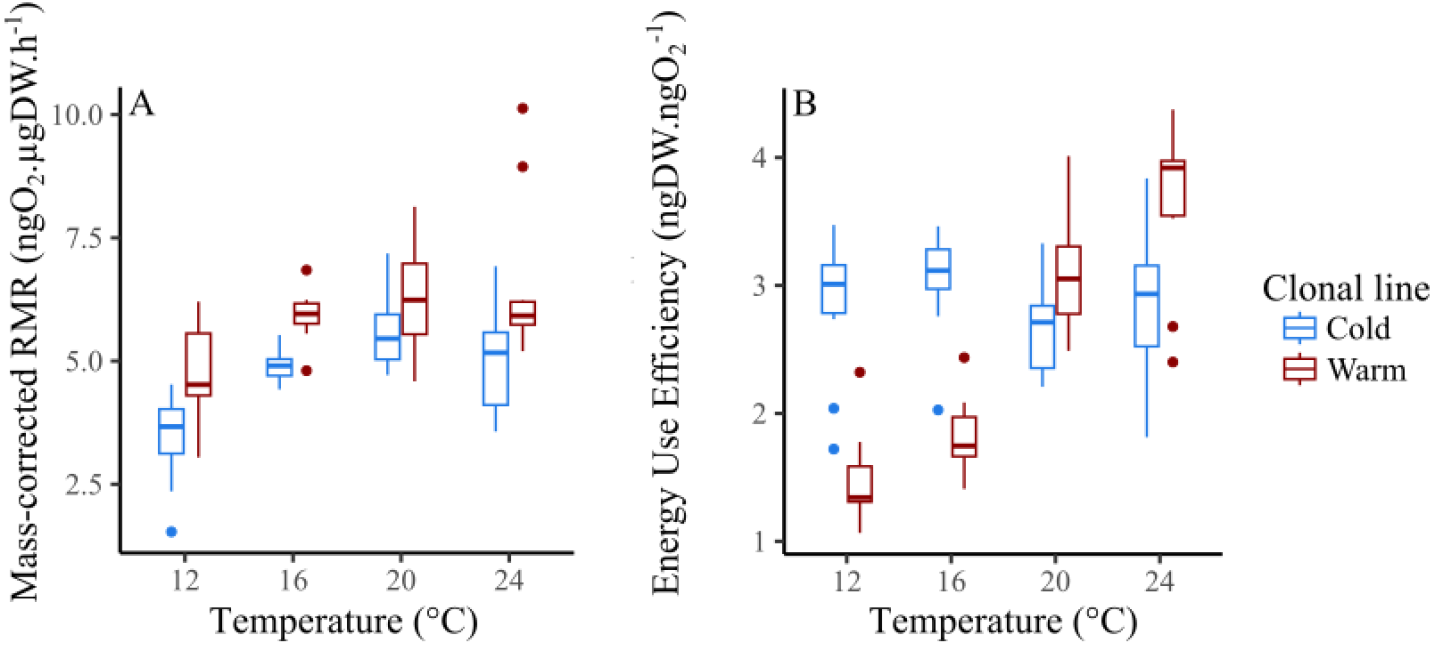
Thermal response of mass-corrected RMR (A) and growth energetic efficiency (B) of the cold and warm clonal line versus thermal gradient. RMR thermal response exhibit a comparable tendency between the cold and warm lines only differing by its overall level. EUE is constant across temperature for the cold line but highly temperature dependent for the warm line.

### Energy use efficiency (EUE)

Our findings revealed distinct thermal responses for the EUE between the clonal lines, as indicated by a significant interaction between temperature and clonal line (ANOVA; F_(1:76)_=65.1; p=9.6e^−12^). The cold line maintained a stable EUE of 3 ng DW ng O□□^1^across the thermal gradient tested (F_(1:39)_=0.63; p=0.43), whereas the warm line exhibited a pronounced temperature-dependent response (F_(1:36)_=109.3; p=2.6e^−12^), with EUE increasing nearly linearly from 1.5 ng DW ng O□□^1^at 12°C up to 3.6 ng DW ng O□□^1^at 24°C. The differential thermal response resulted in higher EUE in the warm line at elevated temperatures, but lower EUE at reduced temperatures, mirroring the thermal growth response (***Fig. 2-B***).

### Model

Our structural model showed that, despite a stronger effect of temperature on individual RMR in the cold line (ω^2^=0.48) versus the warm line (ω^2^=0.25), and a tighter relation between RMR and growth rate (cold ω^2^=0.92; warm ω^2^=0.21), the overall correlation between temperature and growth performance is weaker in the cold (R^2^ = 0.63, RMSE = 0.034) versus warm line (R^2^ = 0.97, RMSE = 0.026). Conversely, the cold line maintained high EUE values regardless of temperature (ω^2^<0.01) versus the strong thermal response of EUE in the warm line (ω^2^: 0.77).

## Discussion

The two clonal lines exhibited a significantly different thermal response, with a thermal optimum 4°C lower for the cold versus the warm clonal line. In parallel, individuals from the cold line also exhibit a larger thermal tolerance breadth than individuals from the warm line. This response pattern highlighted, beside the adaptation to a lower temperature, that the cold line also seemed more generalist than the warm line. This was further supported by the structural model demonstrating that temperature-based predictions of the somatic growth rate were more precise for the warm than for the cold line (***Fig.3***). This difference indicated that the growth performance of the cold line was less tightly coupled to temperature, which matches the conclusion that individuals living near the edge of their thermal niches tended to be thermal specialists whereas individuals in more neutral temperatures displayed a more generalist response (Pörtner 2010; Narum et al. 2013). This phenomenon relates to the “hotter-is-better” hypothesis, arguing that higher temperatures tend to increase the maximal performance of individuals while reducing the range of their thermal tolerance (Angilletta et al. 2010) due to a trade-off between tolerance to high temperature and plastic capacities (Barley et al. 2021). However, the drivers of such generalist-specialist trade-off should be slightly nuanced in our dataset.

**Figure 3.**
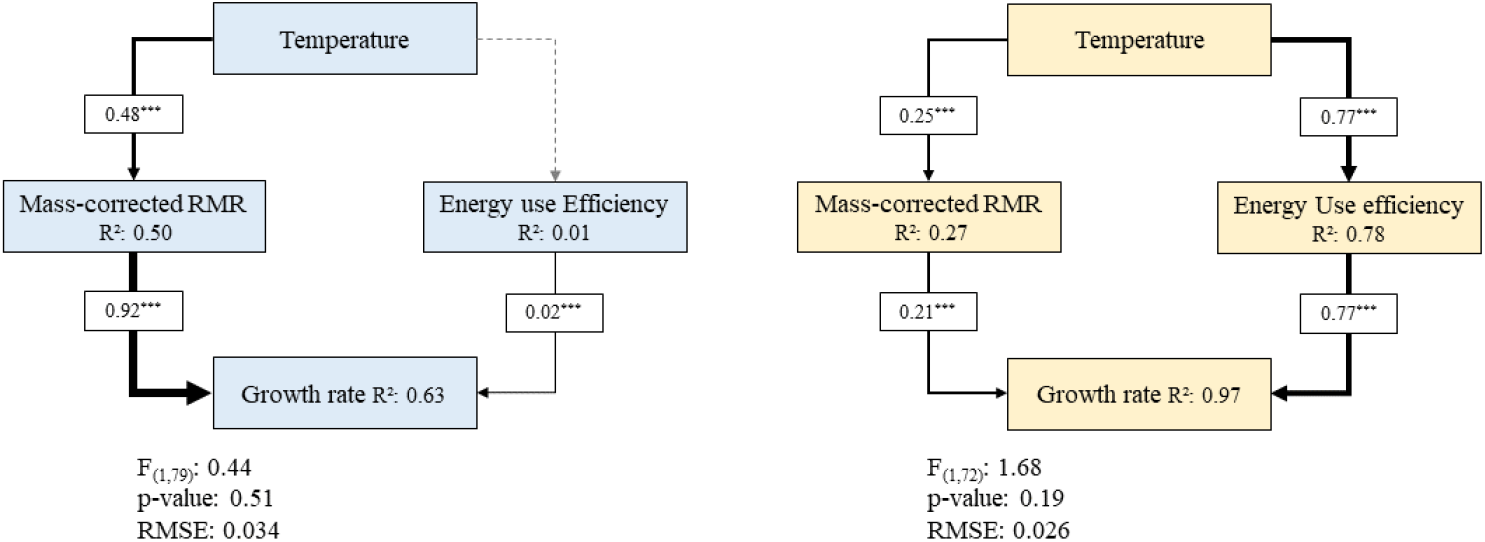
Structural equation model adapted for non-linear relation showing the path effect of temperature on growth with mediating effect of RMR level and EUE. Comparison of the cold (blue) and the warm (yellow) clonal lines. Line thickness reflect the effect strength with values reflecting the effect size (partial omega^2^) and significance levels (0.05*, 0.005**, <0.001***). For SEM models, non-significant p-value for the F-test reflect a good fit.

The cold line was sampled from an aquatic ecosystem with water temperature near 10°C and then kept at 16°C, a process that may have favoured cold generalists over cold specialists. Nonetheless, irrespectively of such a potential sampling bias, the TPCs for somatic growth rates confirm that the cold line was cold-adapted and generalist, whereas the warm line was warm-adapted and specialised. As this work only focus on two specific clonal lines, we should also be careful when generalizing our findings at larger scale. However, given that our primary objective was to elucidate the metabolic pathways and implications of thermal adaptations, rather than their natural selective drivers, using two clonal lines with contrasting thermal responses provide an adequate framework to develop our investigations.

Our results only revealed marginal differences in thermal optimum and thermal breadth of the RMR between the clonal lines. However, we showed that warm-adapted individuals tend to exhibit a mass-specific RMR 30% higher than cold adapted individuals. This trend, while supporting the hypothesis of an adaptive response of RMR to temperature, moves against Krogh’s rule of thermal adaptation of metabolic rate (Ege and Krogh 1914) that argues that warm-adapted individual exhibit lower RMR than cold-adapted ones when exposed to the same temperature (Seebacher et al. 2015; Einum et al. 2019; Jutfelt 2020). Such reduction of metabolic rate in warm adapted organisms is expected to offset the increase of energy and oxygen demand paired with increasing temperature and is sometimes referred to as a metabolic compensation (Havird et al. 2020; Collins et al. 2021). Our results, wherein warm-adapted individuals systematically exhibited higher RMR, challenge this paradigm and support that metabolic compensation may not be a universal feature of thermal acclimation. While exhibiting a similar dynamic than growth rate, the thermal response of RMR here appears only marginal (***Fig.2-A***). While higher inter-individual variability likely contributed to this outcome, this result suggests that the RMR level exhibited a weaker response to temperature than growth, suggesting that RMR might play a less central role in driving thermal adaptation than previously assumed. This observation aligns with emerging questions in thermal biology regarding the role of RMR adaptation in driving ectotherm thermal response (Boltaña et al. 2017; Geisler et al. 2017), an area of research still requiring further investigation.

The weaker role of RMR in mediating individual growth performance across our thermal gradient may be explained by differences in the efficiency of individuals’ growth versus metabolic level (i.e. oxygen consumed). The EUE may constitute a critical point in thermal adaptation and/or acclimation (Pörtner 2010), mediating the relationship between energy requirements (i.e., RMR) and individual growth performance. In our case, we showed that the cold line was able to maintain high level of EUE across the entire thermal gradient, while EUE in the warm lines was strongly mediated by temperature. From a physiological viewpoint, several factors may affect EUE such as differences in enzymatic activities, mitochondrial plasticity or polyunsaturated fatty acid (PUFA) availability. The cell membrane composition, especially in terms of polyunsaturated fatty acids (PUFA) (Müller-Navarra et al. 2004), is known to have strong influences on individual thermal tolerance through homeoviscous adaptation (Masclaux et al. 2009), impacting membrane fluidity and stability (Clamp et al. 1997; Hulbert et al. 2005). Membrane composition also plays a crucial role in influencing individual RMR, as properly functioning membranes facilitate ionic transfer at the cellular level and enhance mitochondrial efficiency (Stanley et al. 2012), thereby increasing the organism’s overall energetic efficiency. In our study, we suggest that the cold-adapted line, which requires higher levels of PUFA to maintain membrane functionality at lower temperatures (Masclaux et al. 2009), displays greater bioconversion capacity to bio-convert PUFA from their dietary precursors. In contrast, warm-adapted organisms, whose environmental PUFA demand is likely reduced, may exhibit lower efficiency in PUFA conversion from their dietary input. Such differences in PUFA bio-conversion capacities may underlie the variation in energetic efficiency observed between the two clonal lines. Cold-adapted lines likely expend less energy in PUFA bioconversion while maintaining a higher PUFA content in their membranes. This optimised PUFA composition supports membrane functionality at low temperatures and enhances the overall energetic efficiency of physiological processes, ultimately improving EUE.

Overall, combining our data and model predictions we showed that despite a stronger effect of temperature on individual RMR in the cold line and a tighter relation between RMR and growth rate, the direct correlation between temperature and growth performance is weaker in the cold versus warm line. This weaker correlation suggests that temperature has a lower effect on individual growth performance for the cold versus warm line, despite its stronger effect on individual RMR. This response here results from the high EUE values maintained by the cold line regardless of temperature versus the strong thermal response of EUE in the warm line. This underscores the critical role of EUE in shaping the thermal response of individual growth performance by mediating the effects of RMR variability. Energy use efficiency may thus constitute a key factor for ectotherms’ thermal adaptation. Moreover, the capacity to maintain a high EUE under a larger thermal gradient can also increase the ability of individuals to tolerate thermal fluctuations, which are expected to increase under global warming. Investigating how EUE stability (or variability) is selected under thermal variation may provide complementary information regarding the evolutionary adaptation of this traits under global warming where both mean temperature and temperature fluctuations are expected to increase.

In conclusion, we demonstrate that, while constituting a main factor mediating the effects of temperature on individual growth performances, RMR appears to be poorly subjected to adaptation between clonal lines with contrasted thermal history. By contrast, we showed that EUE is a critical function mediating the consequences of temperature-driven differences of RMR on individual performances. By increasing their energetic efficiency, individuals were able to offset the inevitable temperature-driven variation of RMR and thus maintaining growth performance over a broader thermal range. Such adaptation to improve individual EUE may pass through maintenance of membrane functionality over a wider thermal range resulting from higher capacities for bio-conversion of membrane building blocks from their dietary precursors. Further studies may scrutinize cell membrane composition dynamics and test whether and how cell membrane PUFA bio-conversion capacities may affect individual RMR and EUE, and ultimately the acclimation capacities of individuals and populations in a context of climate change where temperature, but also dietary supply, are expected to be drastically affected.

## Acknowledgement

This work is part of the project “REBORN” funded under the Esprit program of the Austrian Science Fund (FWF grant N°: ESP 614-B).

