## Supplementary material for "Energy use efficiency may mediate metabolic thermal adaptation in *Daphnia magna*"

| GR | | | |
| --- | --- | --- | --- |
| cold | | warm | |
| maximum | 0.42 [CI-95% 0.33; 0.44] | maximum | 0.55 [CI-95% 0.52; 0.56] |
| optimal T | 19.9°C [CI-95%: 19.5; 20.7]) | optimal T | 23.8°C [CI-95%: 23.1; 24.9] |
| breadth | 10.9°C [CI-95% 9.2; 12.9] | breadth | 8.0°C [CI-95% 7.1; 9.1] |

Supplementary table 1 : Summary of maximum values, optimal temperatures and thermal breadth of juvenile growth rate for the cold and warm lines.

| mcRMR | | | |
| --- | --- | --- | --- |
| cold | | warm | |
| maximum | 7.14 ng O_2_ µg DW h^-1^ [CI-95%: 6.85; 7.55] | maximum | 9.5 ng O_2_ µg DW h^-1^ [CI-95%: 8.9; 11.5] |
| optimal T | 20.24°C [CI-95%: 19.1; 22.65] | optimal T | 23.04°C [CI-95%: 19.4; 44.8] |
| breadth | 16.2°C [CI-95%: 10.4; 19.3] | breadth | 10.9°C [CI-95%: 8.9; 14.4] |

Supplementary table 2 : Summary of maximum values, optimal temperatures and thermal breadth of mass-corrected resting metabolic rate for the cold and warm lines.
